# Electric Field Dynamics in the Brain During Multi-Electrode Transcranial Electric Stimulation

**DOI:** 10.1101/340224

**Authors:** Ivan Alekseichuk, Arnaud Y. Falchier, Gary Linn, Ting Xu, Michael P. Milham, Charles E. Schroeder, Alexander Opitz

## Abstract

Neural oscillations play a crucial role in communication between remote brain areas. Transcranial electric stimulation with alternating currents (TACS) can manipulate these brain oscillations in a non-invasive manner. Of particular interest, TACS protocols using multiple electrodes with phase shifted stimulation currents were developed to alter the connectivity between two or more brain regions. Typically, an increase in coordination between two sites is assumed when they experience an in-phase stimulation and a disorganization through an anti-phase stimulation. However, the underlying biophysics of multi-electrode TACS has not been studied in detail, thus limiting our ability to develop a mechanistic understanding. Here, we leverage direct invasive recordings from two non-human primates during multi-electrode TACS to show that the electric field magnitude and phase depend on the phase of the stimulation currents in a non-linear manner. Further, we report a novel phenomenon of a “traveling wave” stimulation where the location of the electric field maximum changes over the stimulation cycle. Our results provide a basis for a mechanistic understanding of multi-electrode TACS, necessitating the reevaluation of previously published studies, and enable future developments of novel stimulation protocols.

## INTRODUCTION

Efficient brain function relies on a sophisticated multi-scale system of neuronal communication. Neural communication at the level of synaptic signaling gives rise to synchronized excitability fluctuations in neuronal ensembles at the regional and network levels, which in turn emerge as global brain oscillations or rhythms^1^. Brain rhythms reflect neural operations underlying behavior and cognition^2–4^ and are present in all mammals across their evolution^5^. Their abnormality is a significant factor in psychiatric disorders, such as schizophrenia, ADHD, depression and anxiety^6,7^. The rising appeal and evident importance of neural oscillations^8^ that both reflect^1^ and shape^9^ neural communications inspire the search for tools allowing their causal manipulation.

Recent developments enable such manipulations using weak transcranially applied electric fields. Transcranial alternating current stimulation (TACS) can modulate brain rhythms by coupling intrinsic electric fields in the brain to externally applied electric currents in a matching frequency band^10–12^. Such coupling can magnify the power or realign the phase of ongoing brain rhythms. Thus, TACS presents an exciting tool to causally probe the physiological and behavioral role of brain rhythms and their synchronization, i.e., connectivity^13,14^. Several studies have used TACS to manipulate within- and between-area brain connectivity^15–18^. The latter requires a departure from conventional two-electrode stimulation techniques and the use of model-driven, multi-electrode approaches to drive two or more brain sites independently. Through the simultaneous entrainment to the rhythm of external electric fields, researchers can manipulate the phase co-alignment of intrinsic oscillations and study their functional importance.

Modulation of between-area brain connectivity first was done via dual-site TACS where two stimulation electrodes cover two target brain regions and a third electrode serves as a return electrode outside the regions of interest. Following a seminal experimental study^15^, multiple groups employed this procedure^18–21^ or modifications of it^22,23^. So far, all studies have considered two primary stimulation conditions: either the phase of the alternating current was the same (0° shift) between the two stimulation electrodes, or it was the opposite (180° shift). Here, the core idea is that neural oscillations in the target brain areas will resonate with the applied electric currents and align their phases accordingly. Because phase alignment of oscillations in communicating brain regions is functionally important^24–27^, application of in-phase or out-of-phase currents should facilitate or hinder their communication. This principle has guided the design and interpretation of these dual-site and, in general, multi-electrode TACS experiments. However, recent modeling efforts dispute their biophysical validity with regard to the achieved phase differences of stimulation fields^28^.

The current rationale for multi-electrode TACS applications relies on two main assumptions. The first is that the manipulation of phase differences between stimulation electrodes does not significantly change other properties of the generated electric field. Most importantly, this predicts the field’s spatial configuration, i.e., the electric field magnitude at different spatial locations. However, direct measurements to substantiate this assumption are lacking. A second assumption is that phase shifts between stimulation currents translate linearly into phase differences between generated electric fields across locations the brain. This assumption holds for the two electrode case^29^, where the only possible obstacle for such translation could be capacitive effects introducing additional phase shifts. However, phase properties of electric fields arising from multi-electrode TACS have not been considered thus far, either theoretically or experimentally. Here, we aim to address this gap by measuring electric field magnitudes and phase angles across the brain during three-electrode TACS under varying phase conditions. Non-human primate models have proven themselves as useful to study biophysical and physiological TES mechanisms^29–32^ as their brains more closely resemble human brain anatomy than do the brains of non-primate species.

Here, we applied multi-electrode TACS and characterized the spatiotemporal properties of generated electric fields, performing direct, stereotactic EEG measurements (Fig. 1 and supl. animation 1), while varying the phase differences between stimulation currents from 0° to 360° in 15° steps. We found a non-linear relationship between the phase difference of transcranially applied currents and the phases and magnitude of measured intracranial electric fields. We further describe a previously unreported capability of multi-electrode TACS to generate “traveling wave” electric fields in the brain, enabling the design of novel stimulation protocols.

**Figure 1.**
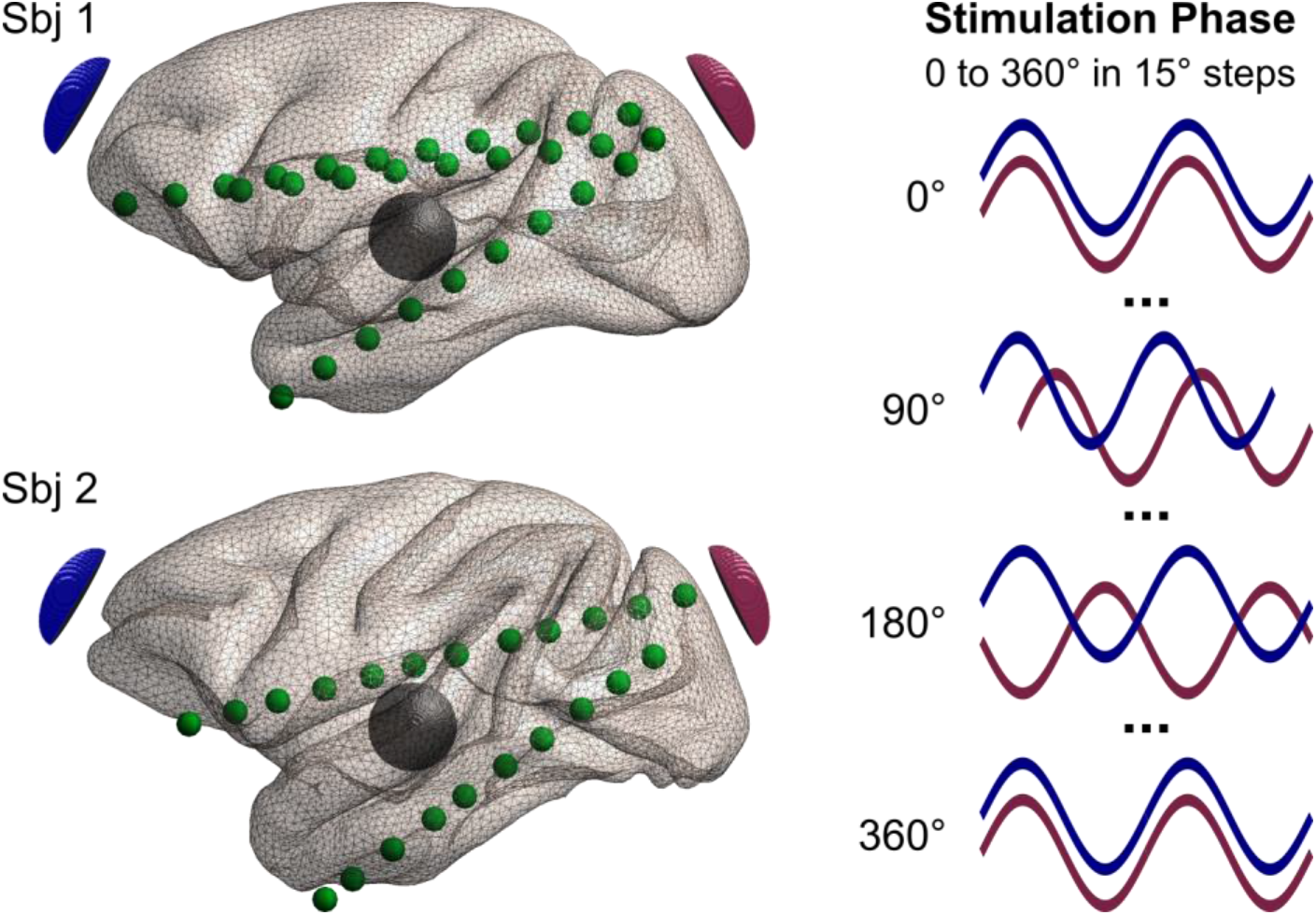
Experimental design. Two non-human primates with recording electrodes implanted along the anterior-posterior plane receive transcranial electric stimulation (TES). For subject 1 we analyze three recording electrodes with 29 contacts altogether, and for subject 2 – two recording electrodes with 22 contacts. Two TES stimulation electrodes are placed on the scalp at anterior (blue) and posterior (red) locations, with the return electrode (black) over the temporal area. Alternating current stimulation was applied at 10Hz and fixed intensity at the stimulation electrodes, while the phase of the stimulation currents between them varied from 0° to 360° in 15° steps (25 phase conditions in total). See supplementary materials for a 3d animation.

## METHODS

### Subjects

Experiments were conducted in two non-human primates. All procedures were approved by the Institutional Animal Care and Use Committee of the Nathan Kline Institute for Psychiatric Research. Subject 1 is a capuchin monkey (*Cebus apella*, 11 y.o., female, 2.9 kg) and subject 2 is a rhesus monkey (*Macaca mulatta*, 6 y.o., female, 4.8 kg). Both subjects were implanted with MRI-compatible head posts and 3 multicontact stereo-EEG depth electrodes (Ad-Tech) that sampled the intracranial electric field at 5 mm intervals in the posterior-anterior direction. The electrodes were implanted via a small craniotomy over the left occipital cortex with the terminal points in the frontal eye field (12 contacts, 5 mm spacing), medial prefrontal cortex (10 contacts, 5 mm spacing) and anterior hippocampus (10 contacts, 5 mm spacing). The craniotomy was sealed after the injection with nonconducting bone cement. The electrode locations were confirmed by post-implantation magnetic resonance imaging.

### Transcranial Alternating Current Stimulation

Transcranial alternating current stimulation (TACS) was delivered using a current-controlled, multi-electrode system StarStim (Neuroelectrics) via three round Ag/AgCl electrodes (3.14 cm^2^) with conductive gel (SigmaGel). The electrodes were placed on the scalp over the middle forehead (first stimulation electrode), the left occipital area (second stimulation electrode) and the left temporal area (return electrode, see Fig. 1). During each experiment, the stimulation intensity (0.1 mA peak-to-zero) and frequency (10 Hz) were kept constant for both stimulation electrodes. The return electrode was set to pick up the remaining currents. The phase of the stimulation currents was varied systematically starting from 0 degrees phase difference between the two stimulation electrodes and up to 360 degrees in 15 degrees steps. This resulted in 25 phase conditions that were measured in both subjects. Each stimulation condition lasted for 30 seconds with a ramp up/down time of 5 seconds.

### Stimulation Current Profile

While the exact amplitude (*A)* and phase (*φ_S_*) of the currents that are passed via the two stimulation electrodes are precisely controlled, the amplitude and phase of the return current are not explicit stated. One can derive the return current analytically as follows:

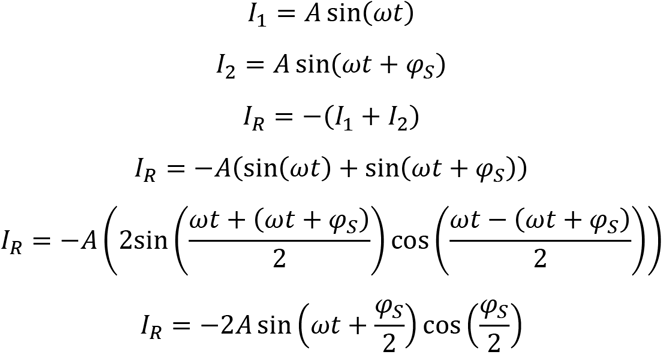

With *I*_1_ – the current of electrode 1 in Amperes, *I*_2_ – the current of electrode 2, *I_R_* – the return current, *t* – the time in seconds, *ω* – the angular frequency in radians per second (*ω* = 2*πf, f* – frequency in Hz), *φ_S_* – the phase difference between the first and second stimulation electrode in radians.

This equation provides a closed form solution for the return current. Amplitude and phase of the return current depend on the phase difference of the stimulation electrodes. For a phase difference of 0 degrees return currents will be maximal, while for 180 degrees return currents are zero. The 180 degrees case corresponds to a classical two electrode TACS configuration. Current profiles for the return current for selected phase conditions are shown in Supplementary Figure 1.

### Intracranial electric field recordings

For subject 1, intracranial electric field recordings were performed using a BrainAmp MR plus amplifier (Brain Products) while for subject 2, a Cortech NeurOne Tesla amplifier (Cortech Solution, Wilmington, NC) was used. Ground and reference electrodes were positioned on the scalp over the right temporal area. Both monkeys were anesthetized (capuchin: ketamine 10 mg/kg IM, atropine 0.045 mg/kg IM, diazepam 1mg/kg IM, isoflurane 2%; macaque: ketamine 12 mg/kg IM, atropine 0.025 mg/kg IM, isoflurane 1.25-2% as needed) during the experiments to reduce the stress of the monkeys and keep them still. While anesthesia suppresses neurophysiological activity, it has negligible impact on electric field measurements which are the objective of this study^29^.

### Data Analysis

Data were analyzed with MATLAB 2017b (MathWorks) according to previously reported processing steps^29,30^. Data preprocessing included bandpass filtering from 5 to 20 Hz with a fourth order forward-reverse Butterworth filter and downsampling to 1 kHz using the Fieldtrip toolbox^33^. Further, we extracted the epochs of interest which correspond to the stimulation conditions (30 seconds per epoch) from the continuous recordings. As a result, 25 epochs were defined for each subject. Finally, we demeaned the data and visually inspected them for the presence of potential technical artifacts. For the subject 1, we excluded three contacts and interpolated them from neighboring sites due to excessive noise at these contacts. We further excluded the most posterior contact for each electrode because they were outside the grey matter. For the subject 2, two contacts were excluded and interpolated from the neighboring sites. One electrode was entirely removed from the analysis due to low recording quality. Finally, we rescaled the data (i.e. multiply by 10) to a 1 mA peak-to-zero stimulation current according to reporting convention for neuromodulation studies^12^. Exemplary data are displayed in Supplementary Figure S2.

After preprocessing, we calculated the amplitude and phase of the recorded TES voltages. The time-series were zero-padded to a length of 32768 (2^15^) samples. Then, we calculated the Fast Fourier Transformation and extracted the phase angles (*φ*) at the stimulation frequency (10 Hz). We unwrapped the phase angles, centered them between −2/π and +2/π, and converted them from radians to degrees. To estimate the maximum phase difference in the brain for a given stimulation condition (Δ*φ*), we subtracted the phase angle from the most posterior recording electrode from the most anterior recording electrode.

In a second analysis step, we calculated the TES-induced electric fields in the brain by computing the numerical gradient of the recorded voltages along the implanted electrodes. Given that the recording electrodes are parallel to the connecting line between two stimulation electrodes, our method accurately captures the dominant anterior-posterior component of the electric field in the brain. Like the TES voltages, the TES electric field is oscillating over time. We estimated the strength of the oscillating electric field as the root mean square amplitude per cycle. This resulted in a more robust estimate compared to the peak amplitude, however with results qualitatively unaffected by this choice. The phase angles (*φ*) of the electric field were calculated in the same way as for the voltage described above.

One intriguing possibility with multi-electrode, multi-phase TACS that we discovered during exploratory data analysis is that the spatial location of maximum electric fields changes with time (i.e., a travelling wave). We thus quantified this travelling wave property as a function of stimulation phase. For this, we first normalized the electric field time-courses for each stimulation condition with respect to their individual maximum. Then, we estimated their pairwise dissimilarity as the mean squared difference across contacts over time and averaged them across electrodes. Dissimilarity close to zero indicates that every recording contact in each electrode detects the highest (or lowest) field magnitudes at the same time point. Thus, the electric field maximum does not move in space over time. High dissimilarity indicates that different contacts detect the electric field maximum at the different time points, i.e., the field’s maximum is moving over time (travelling wave).

### Statistical Analysis

To assess the statistical significance of differences in electric field magnitudes between stimulation conditions we implemented a non-parametric one-way ANOVA test (also known as Kruskal-Wallis test). For measurements where the range of values, rather than the absolute values themselves, are meaningful we utilized a one-way ANOVA that compares the absolute deviations from the group median (Brown-Forsythe test). Such measurements included the maximum voltage and the phase of voltages and electric field. F-statistics, degrees of freedom and p-values are reported in the results section.

Non-linear relationships between the measurement results and stimulation conditions were examined using the MATLAB Curve fitting toolbox. We tested the fit of a linear function (*y* = *a*_1_*x* + *b*) and sinusoidal functions (*y* = *a*_0_ + *a*_1_ cos(*xω*) + *b*_1_ sin(*xω*)). The adjusted R-squared (R^2^_adj_) metrics of goodness-of-fit is reported and complemented by the sum-of-squares-due-to-error (SSE) where appropriate.

### Visualization

All 2d plots were created in MATLAB (MathWorks) and 3d brain plots using Gmsh^34^. For each animal, cortical surfaces were reconstructed in the following steps. First, we employed a bias field correction of the T1-weighted images using ANTs (https://stnava.github.io/ANTs) and averaged them across multiple acquisitions. Brain extraction and tissue segmentation was performed using FreeSurfer (https://surfer.nmr.mgh.harvard.edu) and manual corrections using ITK-SNAP (http://itksnap.org). Finally, the white matter and pial surfaces were reconstructed using FreeSurfer. Figure S3 was drawn with the Gramm toolbox (https://github.com/piermorel/gramm). All figures are assembled in panels in Inkscape (https://inkscape.org). Note that for 10 Hz TES the length of a single cycle is 100 ms which defines the time window for the corresponding figures and animations.

## RESULTS

To assess the voltage and electric field distributions in the brain arising from multi-phase TACS, we conducted direct, intracranial measurements in two non-human primates (Fig. 1 and supl. animation 1).

The voltage distribution in the brain demonstrates a clear dependence of the stimulation phase (Fig. 2A, B). A much smaller range of TES voltages in the brain was found for the 0° condition with 6.64 mV for the subject 1 and 15.19 mV for the subject 2, compared to the 180° condition with 30.18 mV and 43.77 mV respectively. One-way (factor: applied stimulation phase) ANOVA of deviations from the median confirms a strong disparity in voltages across the stimulation conditions (subject 1: F_24,700_ = 15.54, p = 3.3×10^−50^; subject 2: F_24,525_ = 7.45, p = 1.6×10^−21^). The TES voltage gradient is oriented along the anterior-posterior plane with its minimum and maximum occurring in the proximity to the target stimulation electrodes (Fig. 2C, D).

**Figure 2.**
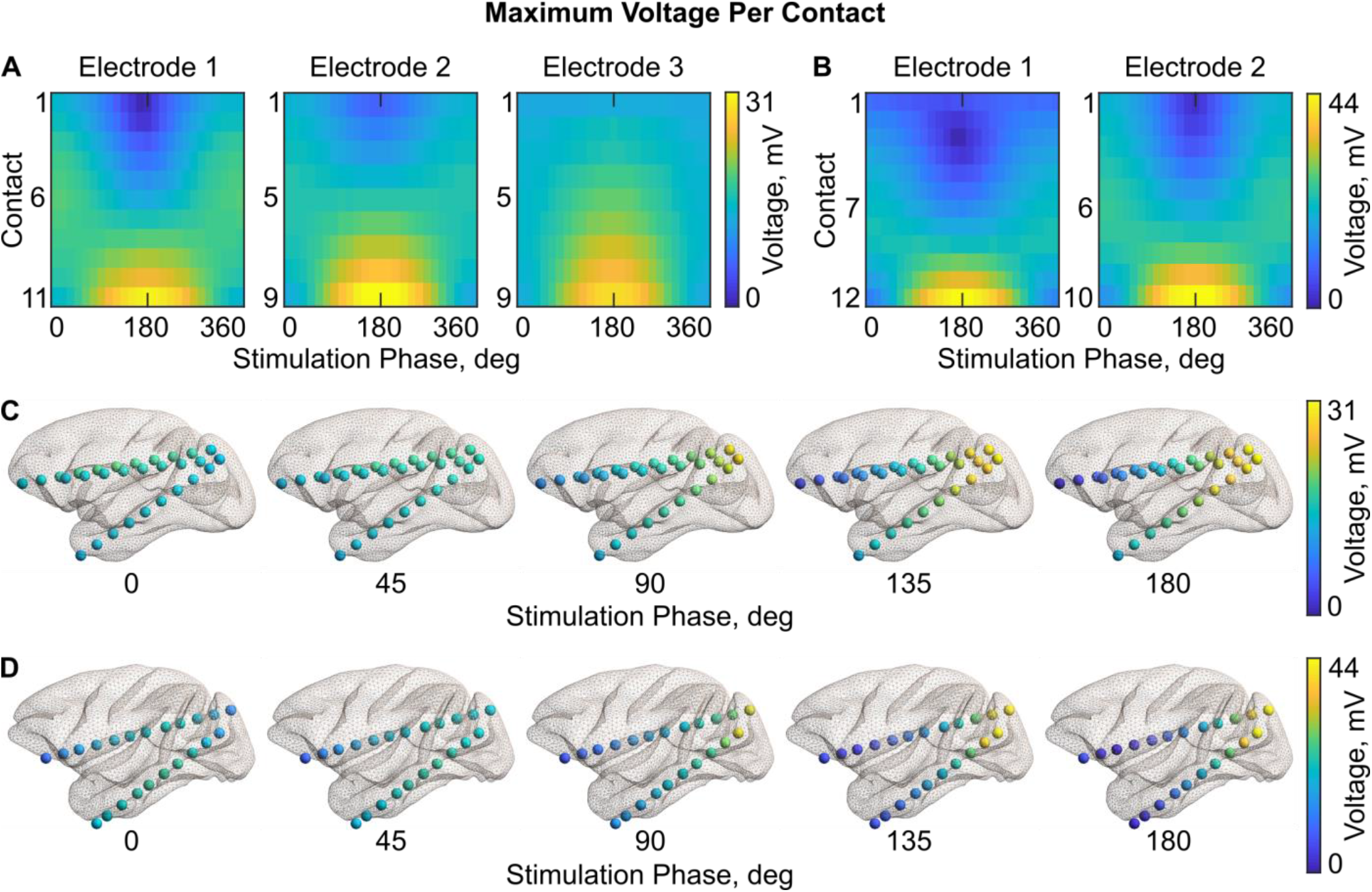
Voltage distribution during TES for different stimulation conditions. (A) Heatmaps of the maximum voltage for each recording electrode for subject 1 (left) and subject 2 (right, B). The x-axis indicates the applied phase difference between the anterior and posterior stimulation electrodes from 0° to 360° in 15° steps, and the y-axis corresponds to the recording contacts (from the first – most anterior contact, to the last – most posterior contact). (C) Voltage gradients for select stimulation conditions for subject 1 and subject 2 (D) visualized at their recorded anatomical locations.

We found a similar picture for the magnitude of the electric field (Fig. 3A, B Fig. S3). The maximum electric field is significantly weaker for the 0° than for 180° condition (non-parametric one-way ANOVA for the subject 1: χ^2^_24,700_ = 396.2, p = 4.5×10^−69^; for the subject 2: χ^2^_24,525_ = 164.3, p = 6.95×10^−23^). Relatively higher electric field magnitudes in posterior brain regions are due to their closer proximity to the stimulation electrode compared to anterior brain regions. Further, we found the electric field strength to be higher in superficial brain regions for the 0° condition and higher magnitudes at progressively deeper brain regions with increasing stimulation phase difference up to 180°. (Fig. 3C, D). The relative ratio of the electric field strength at the most superficial contact compared to the deepest recording contact were found as follows: subject 1 at 0° condition = 11.35, at 45° = 2.59, at 90° = 2.09, at 180° = 1.93; for subject 2 at 0° = 10.95, at 45° = 3.32, at 90° = 2.79, at 180° = 2.5.

**Figure 3.**
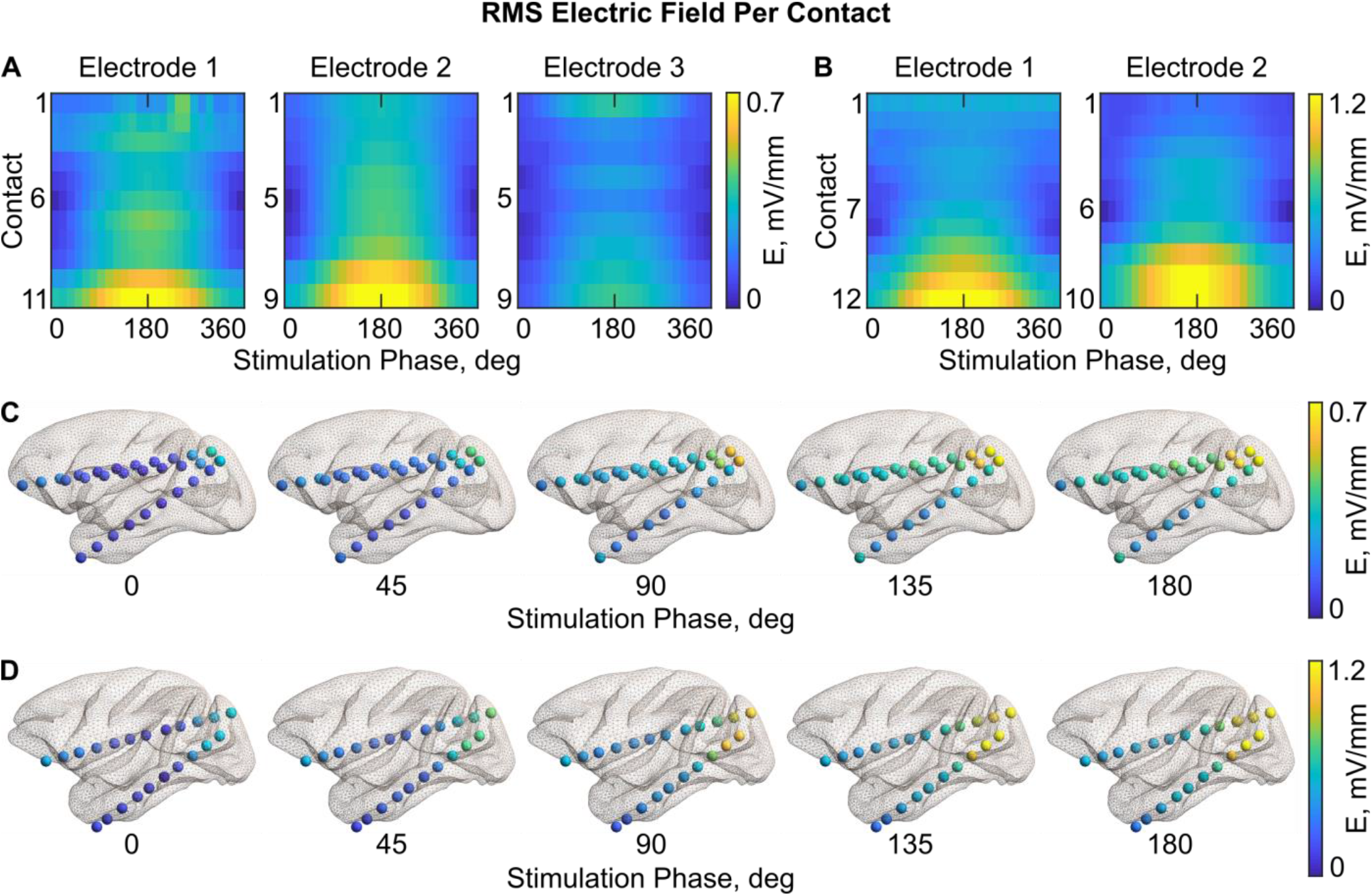
TES electric field distribution for different stimulation conditions. (A) Heatmaps of the root mean square magnitude (RMS) of the electric field for each recording electrode for subject 1 (left) and subject 2 (right, B). The x-axis indicates the applied phase difference between the anterior and posterior stimulation electrodes from 0° to 360° in 15° steps, and the y-axis corresponds to the recording contacts (from the first – most anterior contact, to the last – most posterior contact). (C) Electric field distributions in the brain for select stimulation conditions for subject 1 and subject 2 (D). See also supplementary figure 3 for further details.

Further investigating the stimulation phase dependency of TES voltages and electric fields, we observed a non-linear increase from 0° to 180° and a non-linear decrease from 180° to 360°. We found that a sinusoidal curve fit the voltage data very well, SSE (sum of squares due to error) = 1.29 and R^2^_adj_ = 0.99 for the subject 1, and SSE = 4.25 and R^2^_adj_ = 0.99 for the subject 2. For comparison, linear fit exhibits an order of magnitude larger errors (for the subject 1: SSE = 29.9, R^2^_adj_ = 0.91; for the subject 2: SSE = 71.45, R^2^_adj_ = 0.91). A similar relationship was observed for the magnitude of the electric field. A sinusoidal curve fit the data almost perfectly (for subject 1: SSE = 0.001, R^2^_adj_ = 0.99; for subject 2: SSE = 0.001, R^2^_adj_ = 0.99), though the linear fit is nearly as good (for subject 1: SSE = 0.01, R^2^_adj_ = 0.96; for subject 2: SSE = 0.04, R^2^_adj_ = 0.95).

Next, we analyzed how the phase angles of voltage (*φ_V_*) and electric field (*φ_E_*) in the brain depend on the stimulation conditions. The conditions strongly affect *φ_V_* (One-way ANOVA of deviations from the median for subject 1: F_24,700_ = 6.38, p = 1.76×10^−18^; subject 2: F_24,525_ = 4.54, p = 2.02×10^−11^) and *φ_E_* (subject 1: F_24,700_ = 40.16, p = 1.85×10^−114^; subject 2: F_24,525_ = 9.73, p = 3.83×10^−29^) although in different ways. To analyze how large phase differences can occur across the brain, we computed the phase difference between the most anterior and most posterior recording contact for each electrode. For the voltage phase (Δ*φ_V_*), we found close to zero differences for both 0° (equivalent to 360°) and 180° stimulation conditions (Fig. 4A, C, D) as previously reported^29^. Δ*φ_V_* is maximal for 135°-150° conditions and, symmetrically, for 210°-225° conditions. At the same time, the anterior-posterior difference in phase angles of the electric field (Δ*φ_E_*) is maximal for 0°/360° conditions and close to zero for 180° condition (Fig. 4B). Once again, the changes are not linear across conditions, and are better approximated by a sinusoidal curve (all R^2^_adj_ ≥ 0.98). Considering the spatial distribution of Δ*φ_E_* (Fig. 4E, F; and Fig. S4, S5), the zero-phase difference during 180° stimulation condition indicates a unidirectional electric field along the recording electrodes. Large Δ*φ_E_* characteristic for the 0° stimulation condition is a sign of anti-directional electric fields, such as inward- or outward-oriented.

**Figure 4.**
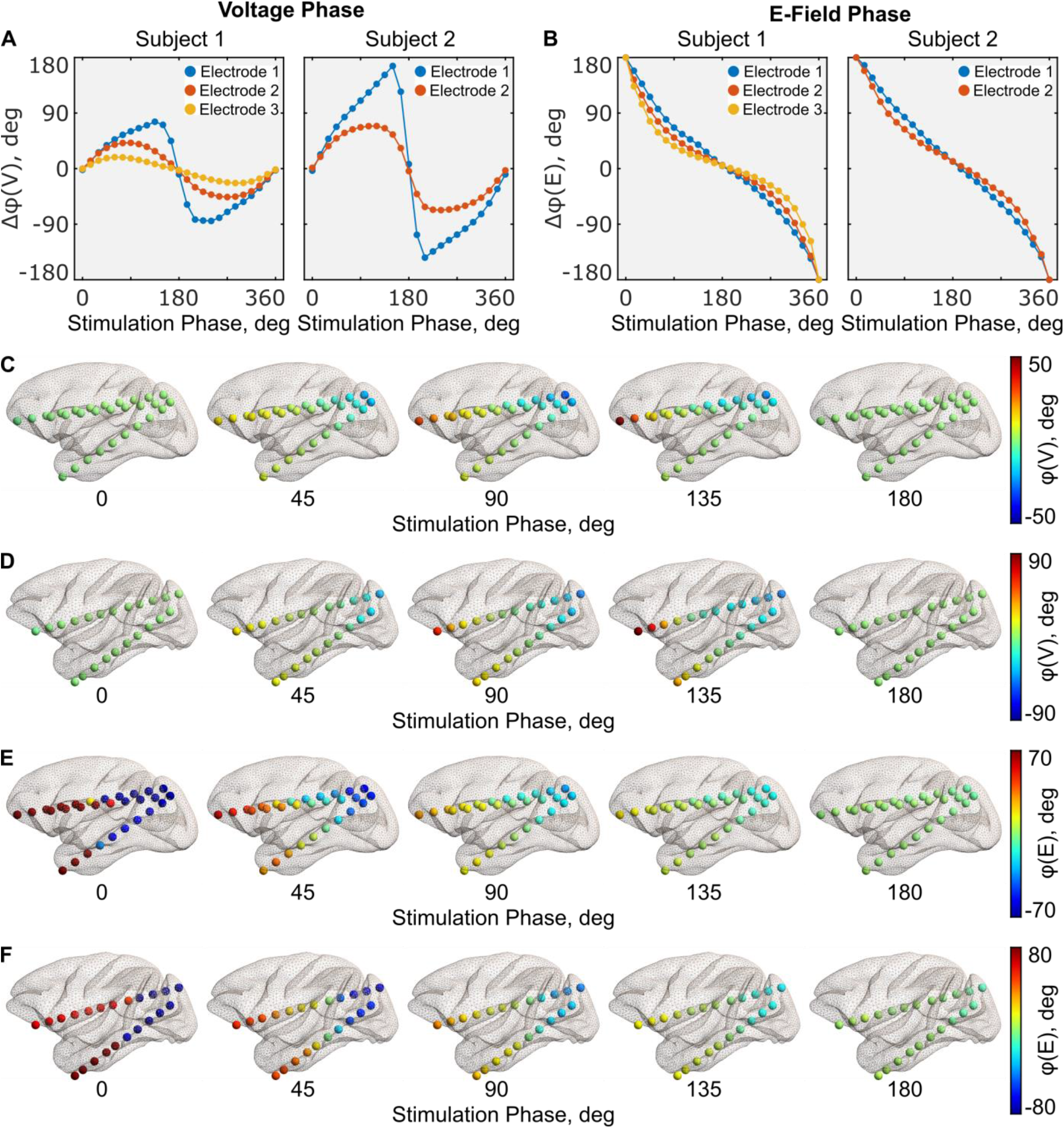
Voltage and electric field phases in the brain during TES for a given stimulation condition. (A) The difference between the voltage phases recorded from the most anterior and most posterior contact (Δ*φ_V_*) per each electrode. The left figure corresponds to subject 1, and the right figure to subject 2. (B) Same in panel A, but for the electric field phase differences (Δ*φ_E_*). (C) 3D visualizations of the voltage phases (*φ_V_*) in the brain for the primary stimulation conditions at all recording contacts for subject 1 and subject 2 (D). (E-F) Same as in panels C and D, but for the electric field phase (*φ_E_*). See also the supplementary figures 4 and 5 for further detail.

In further analysis, we investigated the evolution of electric fields over time. Electric fields in the brain vary over time with the same period as the applied current (Fig. S2). Figure 5 shows the temporal evolution of electric fields for three main stimulation conditions (0, 90 and 180 degrees, see also supl. animation 2, 3). Stimulation in the 180° condition leads to a unidirectional electric field for all time-points. The electric field direction is periodically switching from an anterior->posterior to a posterior->anterior orientation. Stimulation in the 0° condition has a reverse effect: the electric field is anti-directional between the anterior and posterior brain region for all time-points while the direction is changing between inward and outward orientations. Stimulation in the 90° condition creates an intermediate scenario with the electric field being unidirectional for 50% of the time, inward-oriented for 25% and outward-oriented in another 25% of the time. The spatial location of the maximum of the electric field strength was found to be at the same location for the 0° and 180° stimulation conditions for all time points. Intriguingly, intermediate stimulation conditions, such as the 90° condition, can generate a “traveling wave” in the location of the electric field maximum. This means that the spatial location of the stimulation maximum at a given time point varies over the stimulation period (supl. animation 4).

**Figure 5.**
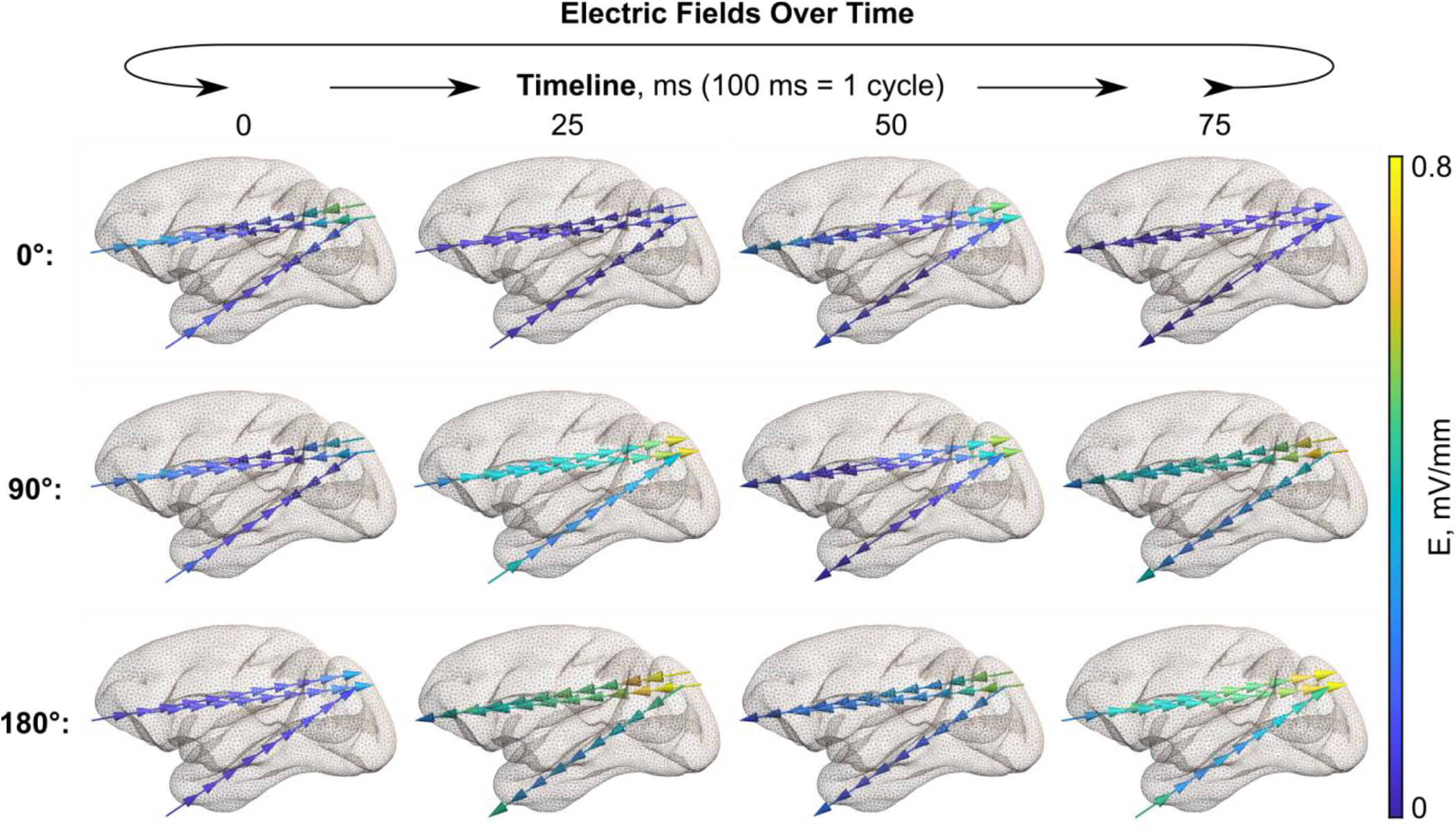
Electric fields in the brain over time during TES for a given stimulation condition. The panel depicts the main conditions (0°, 90°, and 180° stimulation phase differences) for subject 1. Arrows indicate the electric field direction, and the color encodes the electric field magnitude. See supplementary figure 6 for subject 2 and supplementary 3d animation for more details.

In an additional analysis we further quantified this “traveling wave” phenomenon in more detail. For this we normalized the electric field time-courses to the individual maximum for each stimulation condition and each recording contact. This was done such that different electric field distributions can be easily compared across all stimulation conditions and locations. We defined a dissimilarity index as the mean squared difference of the normalized electric fields across time points. A dissimilarity index score closes to zero was found for the 0° and 180° stimulation conditions, indicating that for each contact the electric field maximum occurred at the same time point (Fig. 6A, B). However, other stimulation conditions, most noticeably the 45° condition and symmetric to it the 315° condition, are characterized by a strong dissimilarity of electric field time-courses. This means that the spatial locations of their electric field maxima are traveling over time. Further, the spatial location of these electric field maxima showed a clear temporal pattern (Fig. 6C and supl. animation 4). We found that the relative maximum of the electric field is periodically and gradually moving between the most anterior and the most posterior contact, thus creating a “traveling wave” of electric stimulation.

**Figure 6.**
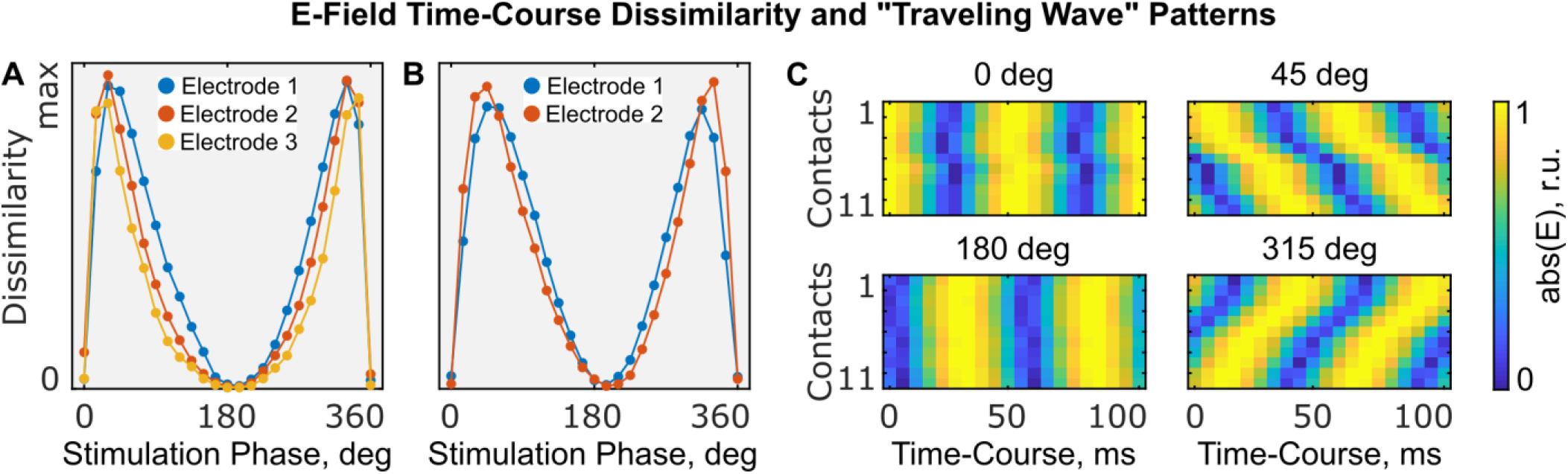
Traveling wave stimulation. Electric field time-courses across different recording contacts (= anatomical locations) are identical for 0°/360° and 180° stimulation conditions but demonstrate traveling wave properties for intermediate stimulation conditions e.g. 45°. (A+B) Dissimilarity (mean square difference) between the electric field time-courses at different contacts. The left figure corresponds to subject 1, and the right figure to subject 2. (C) Absolute normalized (per location) electric field time-courses for subject 1 and electrode 1 in the relative units (r.u.). The first contact in the electrode corresponds to the most anterior location, and the last contact – to the most posterior location. While for 0° and 180° the maxima across contacts occur at the same time point, they occur at different time points for the 45° condition (= traveling wave). Other electrodes demonstrate a similar pattern. See supplementary materials for a 3d animation.

## DISCUSSION

The present work quantified the biophysical features of multi-electrode, multi-phase TES, which forms the foundation for explaining and predicting its neuromodulatory effects. We directly recorded and analyzed the electric potential and field in the non-human primate brain arising from three-electrode transcranial alternating current stimulation. Our study has three main findings: (i) differing electric field magnitude across stimulation phase conditions; (ii) non-linear relationship between transcranial stimulation phase and measured intracranial phase; (iii) specific phase configurations can create travelling wave stimulation patterns. Importantly, all measurements were repeated in two non-human primates from two taxonomic families with substantially different head anatomy. Our clear and highly comparable results encourage the generalization and translation of the present findings to all primates, including humans.

The dominant method of dual-site TACS in human studies is the application of a three-electrode montage where two stimulation electrodes are either in-phase (0° condition) or in anti-phase (180° condition) with each other^15,18–21^. One key assumption behind this approach is that the generated electric field is comparable in any aspect but phase. However, recent modelling work challenged this assumption and demonstrated that electric fields differ significantly between the two conditions^28^. Our experimental data support and extend these model-driven considerations. We found that the electric field strength is significant larger (2-2.3 times) for the anti-phase than for the in-phase stimulation condition. This difference in electric field strength should thus be revisited as a potential driver of the behavioral or physiological effects reported between in-phase or anti-phase stimulation in the literature. This will also apply to four- and more electrode montages where each brain area of interest is covered by a single electrode^22,23^. Importantly, the electric field magnitude between stimulation conditions is changing in a non-linear manner which is well approximated by a sinusoidal curve. Here, it is important to note that the measured electric fields are restricted to their anterior-posterior component along the implanted electrodes. It is to be expected that in the 0° conditions medial-lateral components will be larger than for the 180° condition. This does not diminish the validity of our findings as the direction of the electric field with respect to neural elements is most important for TES physiological effects^35,36^. Thus, differences in one electric field component across stimulation conditions are important to document and control for. Here, our measurements can inform future experimental work so that it can better control for these effects.

Besides the electric field magnitude, we analyzed the electric field phase as a function of stimulation condition. Contrary to the intuitive expectation, the in-phase (0°) stimulation condition resulted in an “antiphase” electric field, where anterior and posterior brain areas experience anti-directional electric fields at any moment in time. The spatial location of the deflection point (from anterior oriented to posterior oriented electric fields) crucially depends on the location of the return electrode and future studies can aim to manipulate this by carefully choosing the location of stimulation and return electrodes. The antiphase (180°) condition leads to an “in-phase” or unidirectional electric field for all contacts and all time-points. Here, the electric field direction is flipping over time between anterior- and posterior-directed states. These differences in electric field directionality might be a driving force for previous findings. Of course, this is complicated by the fact of differing neuronal orientations with respect to the TES electric field in a folded cortex^37^.

The picture for intermediate phase conditions, such as 90° or 270°, is more intricate. It is a mixture of anti- and uni-directional electric field states that reoccur over time. Overall, the complexity of the relationship between phase conditions in multi-electrode TES and electric field phase presents a challenge and an opportunity for future experimental and modeling studies. Precise measurements are especially important for efforts regarding TES artifact rejections due their interaction with physiological signals^38^. Our experimental data underscore the point that explicitly accounting for the full temporal dynamics of electric fields can significantly expand the understanding of TES effects, especially for multi-electrode stimulation.

One novel, intriguing finding that arises from the temporal analysis of TES electric fields across the whole spectrum of phase conditions is a “traveling wave” stimulation condition. It is characterized by a gradual and periodic movement of the electric field maximum in the brain during multi-electrode, multi-phase alternating current stimulation. Such propagation of electric fields resembles the well-known electrophysiological phenomenon of traveling waves in the brain^39,40^. At the macroscopic level, these traveling waves of electrical activity contribute to the formation of large-scale cell assembles and neural networks. Traveling waves are observed and functionally relevant in different frequency bands, brain states and anatomical regions, including theta activity in the hippocampus^41^, beta activity in the motor cortex^42^ and gamma activity in the visual cortex^43^. Traveling waves can even occur on a global level spanning the whole neocortex^44,45^. Thus, mimicking and manipulating such naturally occurring brain activity with TES presents a novel attractive stimulation possibility. New computational tools to optimize electrode placement and timing of input currents will be needed for a successful practical implementation.

The present study confirms and expands previous work on the biophysical mechanisms of TES in non-human primates^29,30^ and epilepsy patients^46^. Previous studies conducted for classic two-electrode montages found that TACS generated electric fields behave in an ohmic manner with minor-to-negligible phase shifts in the low frequency range. We can thus generalize our results to other low frequencies in the EEG range as well. Using phase-shifted input currents from a three-electrode montage, we demonstrate the possibility of phase differences of TES voltages and electric fields. This opens the possibility to create stimulation protocols to stimulate remote brain regions at different phase relations. While this was intended in previous studies^15,18,19,21–23^, they were not based on biophysical considerations of TES electric fields. Future work that includes new optimization schemes in modelling packages such as SimNIBS^47^ will allow a more principled design of multi-phase stimulation protocols.

In summary, the findings of the present work address important biophysical aspects of multi-electrode TES. Previous studies investigated the effect of electric fields on the neurophysiology using various experimental approaches from *in silico*^48^ to *in vitro*^49–51^ and *in vivo*^52^. Seminal work from Terzolo (1956) demonstrated the sensitivity of neurons to weak electric fields (≈1mV/mm) strongly depending on the electric field orientation with respect to the somatodendritic axis^49^. More recent studies show the importance of cellular and network level states, and their coherence with the stimulation parameters^48,50–52^. However, surprisingly little is known of the present electric fields during TES in the brain. System level effects of TES are driven by physical factors such as “where” and “when” the generated electric fields interact with neural tissue. While the “where” question has been explored in previous studies^53–55^, the “when” question specific for alternating current stimulation was largely missing thus far. Here, we examined the spatiotemporal properties of multi-electrode TACS with direct, stereotactic recordings in the non-human primate brain. The results show the importance of the stimulation phase both for the phase and magnitude of the generated electric field. Further, we demonstrated that multi-electrode, multi-phase TES can create a “traveling wave” stimulation where the location of the maximum stimulation changes over time. The present study opens new possibilities for exploiting the temporal complexity of multi-electrode, multi-phase TES and enables future efforts to tie together the “when”, “where” and “how” of TES into a comprehensive, predictive model.

## ACKNOWLEDGMENTS

Research reported in this publication was supported by NIH (R21 MH110217-01, R01 MH 111439 1 and P50 MH109429) and the University of Minnesota’s MnDRIVE Initiative. We thank Stan Colcombe, Raj Sangoi and Caixia Hu for MR imaging support, and we thank Deborah Ross, Mark Klinger, Kathleen Shannon and Tammy McGinnis for veterinary assistance.

## AUTHOR CONTRIBUTIONS

A.O. designed the experiments; A.Y.F., G.L. and A.O. collected the data; T.X. processed the MR data; C.E.S. and M.P.M supervised the data collection; I.A. analyzed the data and prepared the figures; A.O. supervised the data analysis; I.A. and A.O. interpreted the results; I.A. and A.O. wrote the manuscript; all authors reviewed the manuscript.

## SUPPLEMENTARY MATERIALS

**Supplementary Figure 1.**
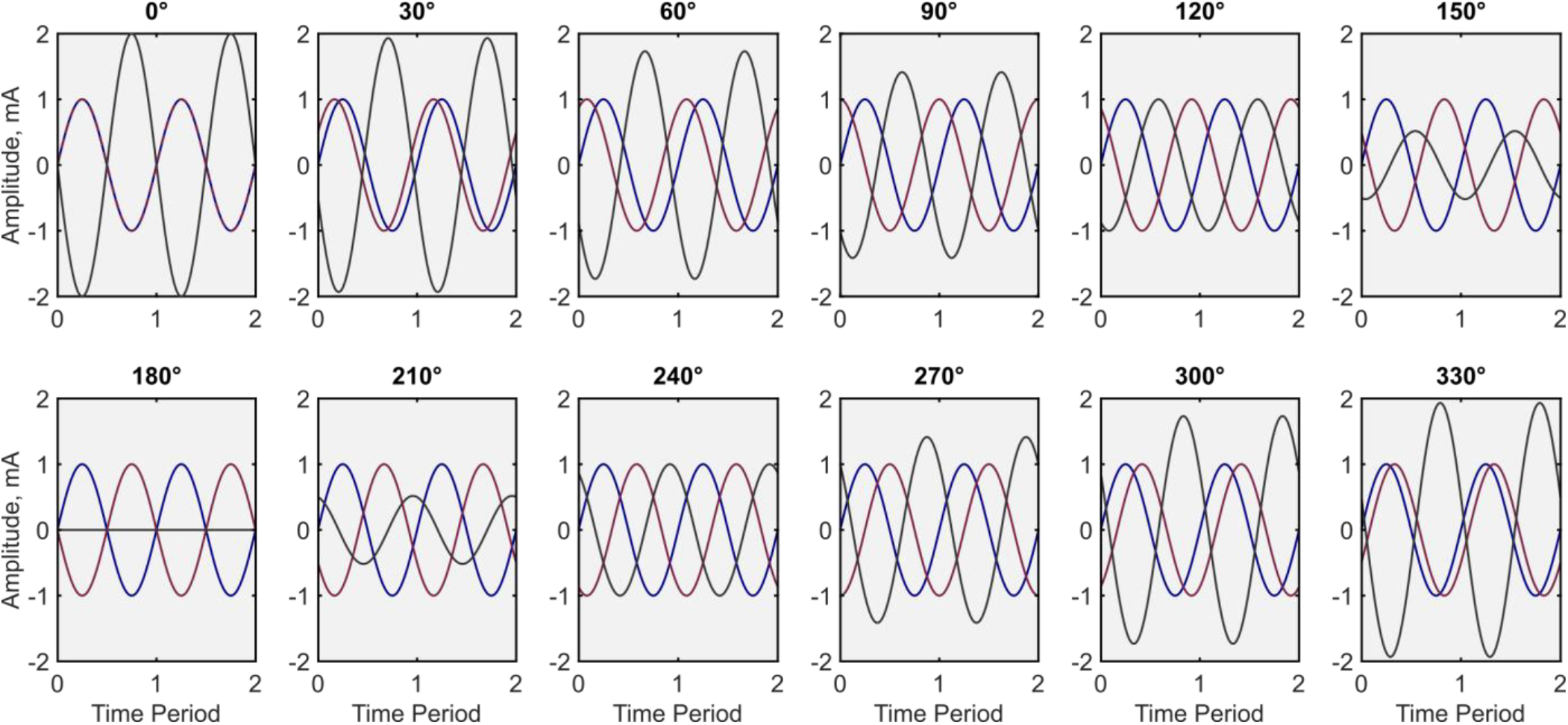
Electric currents that are passed through the first stimulation (blue), second stimulation (red), and return (gray) stimulation electrodes depending on the stimulation condition. Stimulation conditions vary in the phase difference between the currents that pass via the two stimulation electrodes. In this illustration, the peak-to-zero intensities of stimulation currents are constant and equal to 1 mA. This figure complements the chapter “Stimulation Current Profile” in the methods section.

**Supplementary Figure 2.**
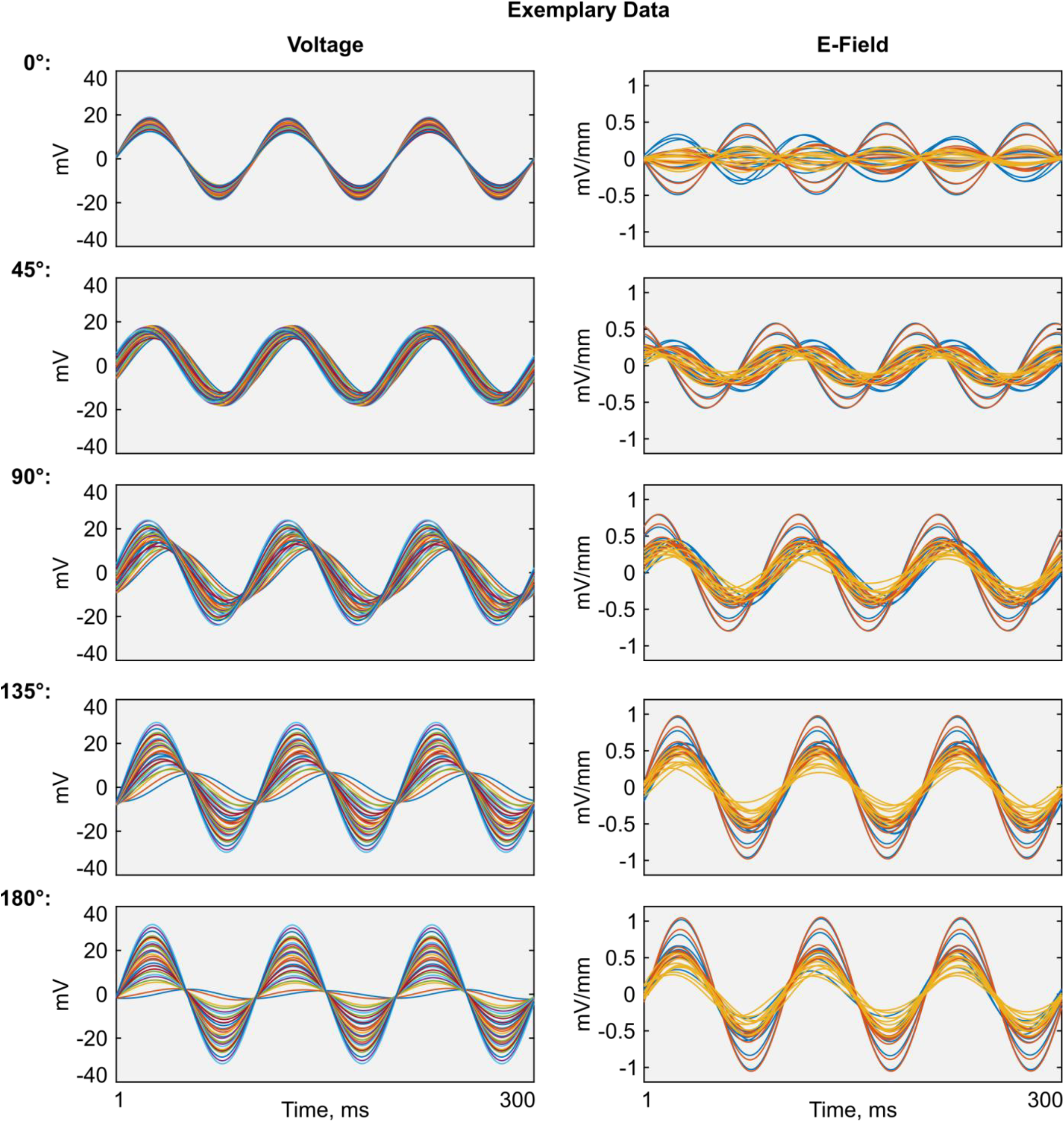
Examples of preprocessed recordings from the implanted electrodes during TES for five stimulation conditions (from top to bottom: 0°, 45°, 90°, 135° and 180° stimulation phase differences between the anterior and posterior stimulation electrodes). Here and everywhere, the units of the raw data are scaled to mV (for the voltages) or mV/mm (for the electric field) per 1 mA peak-to-zero of transcranially applied electric current.

**Supplementary Figure 3.**
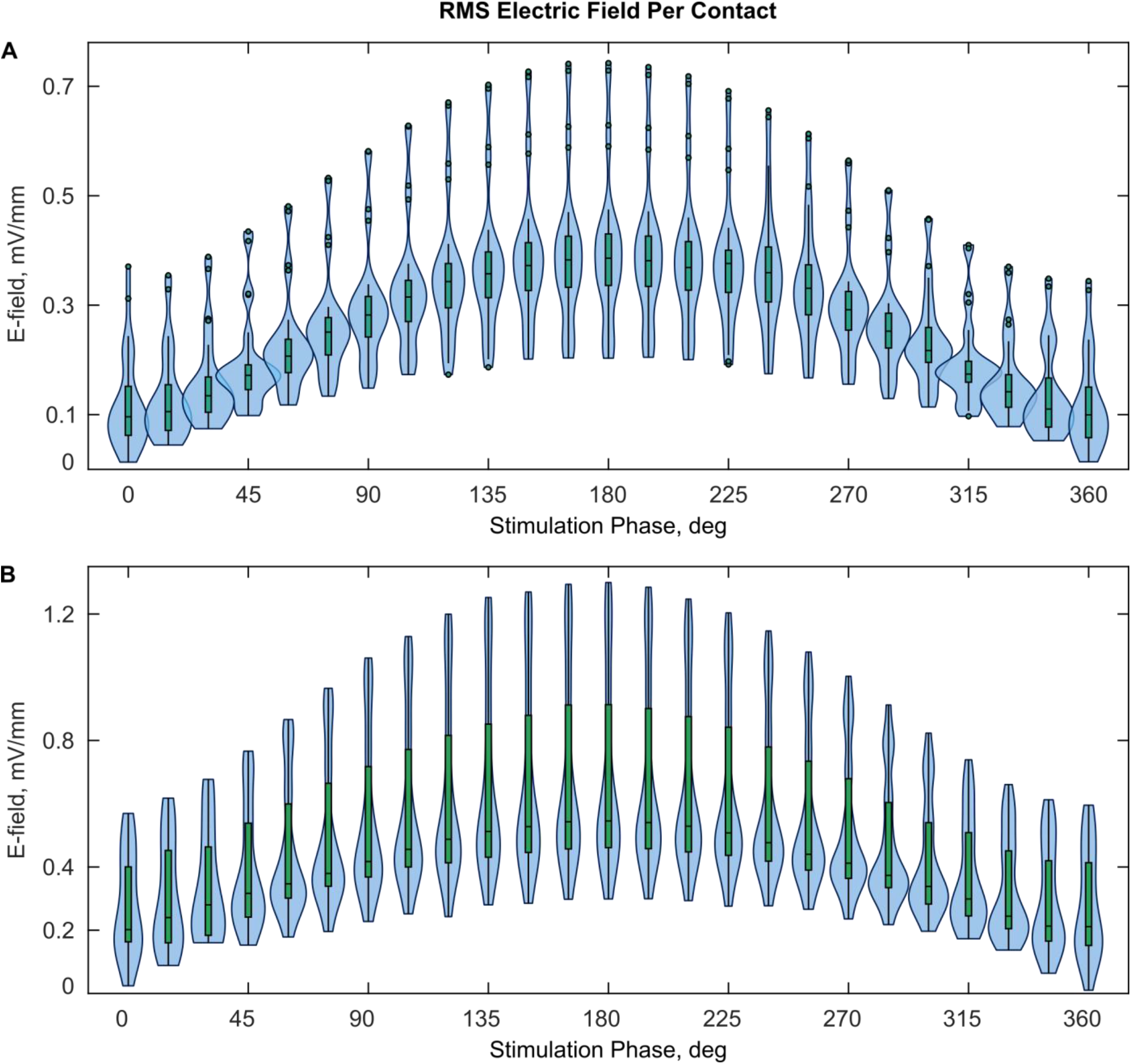
Violin plots of the RMS magnitude of the electric field in the brain. The x-axis indicates the stimulation condition (i.e., the phase difference between the anterior and posterior stimulation electrodes). The upper plot (A) corresponds to subject 1, and the lower plot (B) to subject 2. The kernel density outlines are normalized to have the same area. The units are scaled to mV/mm per 1 mA peak-to-zero of applied current. This figure complements Figure 3 in the main paper.

**Supplementary Figure 4.**
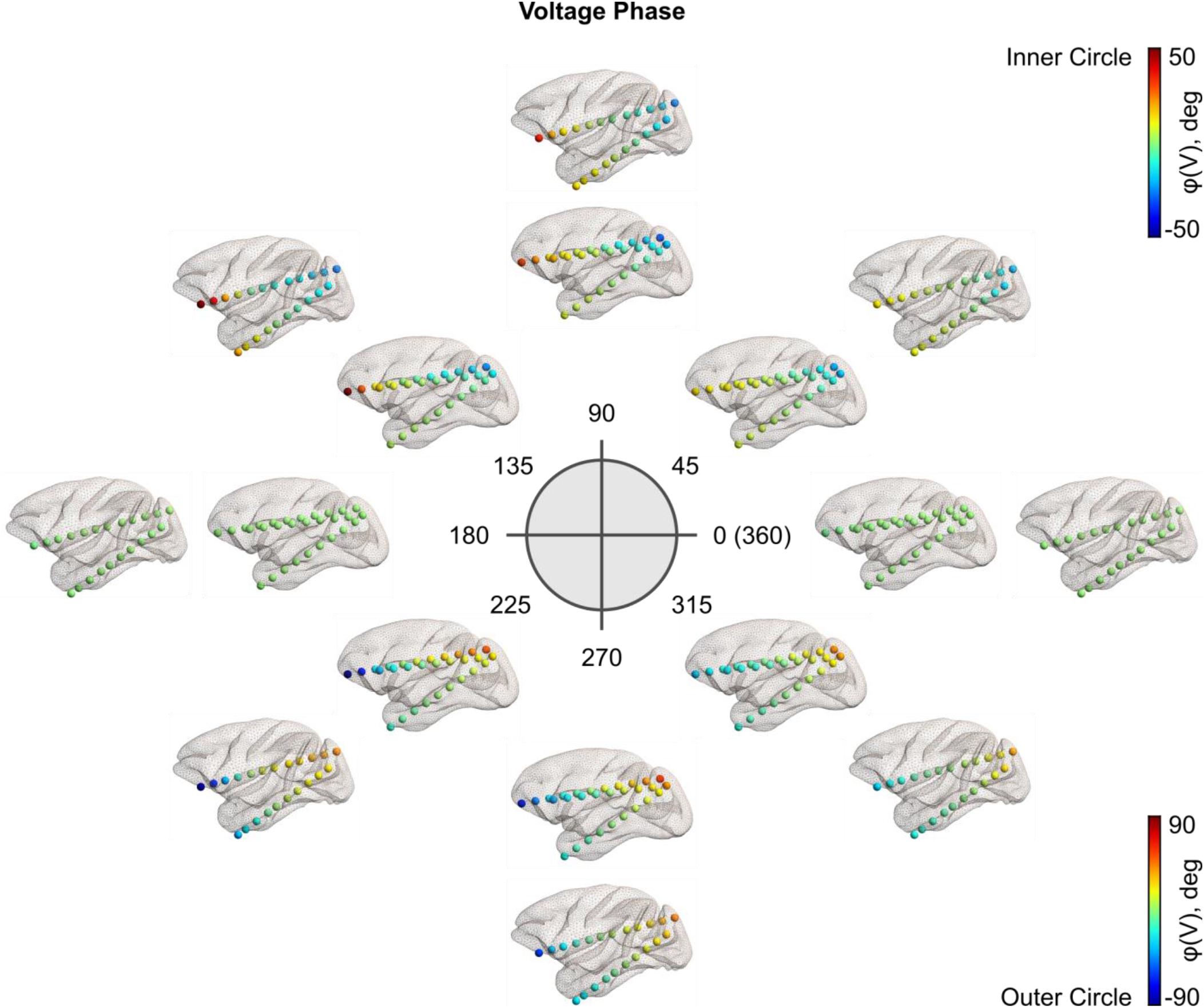
Voltage phases in the brain during TES. The phase difference between the anterior and posterior stimulation electrodes is indicated in the middle plot. The inner circle of plot corresponds to subject 1, and the outer circle to subject 2. This figure complements Figure 4 in the main paper.

**Supplementary Figure 5.**
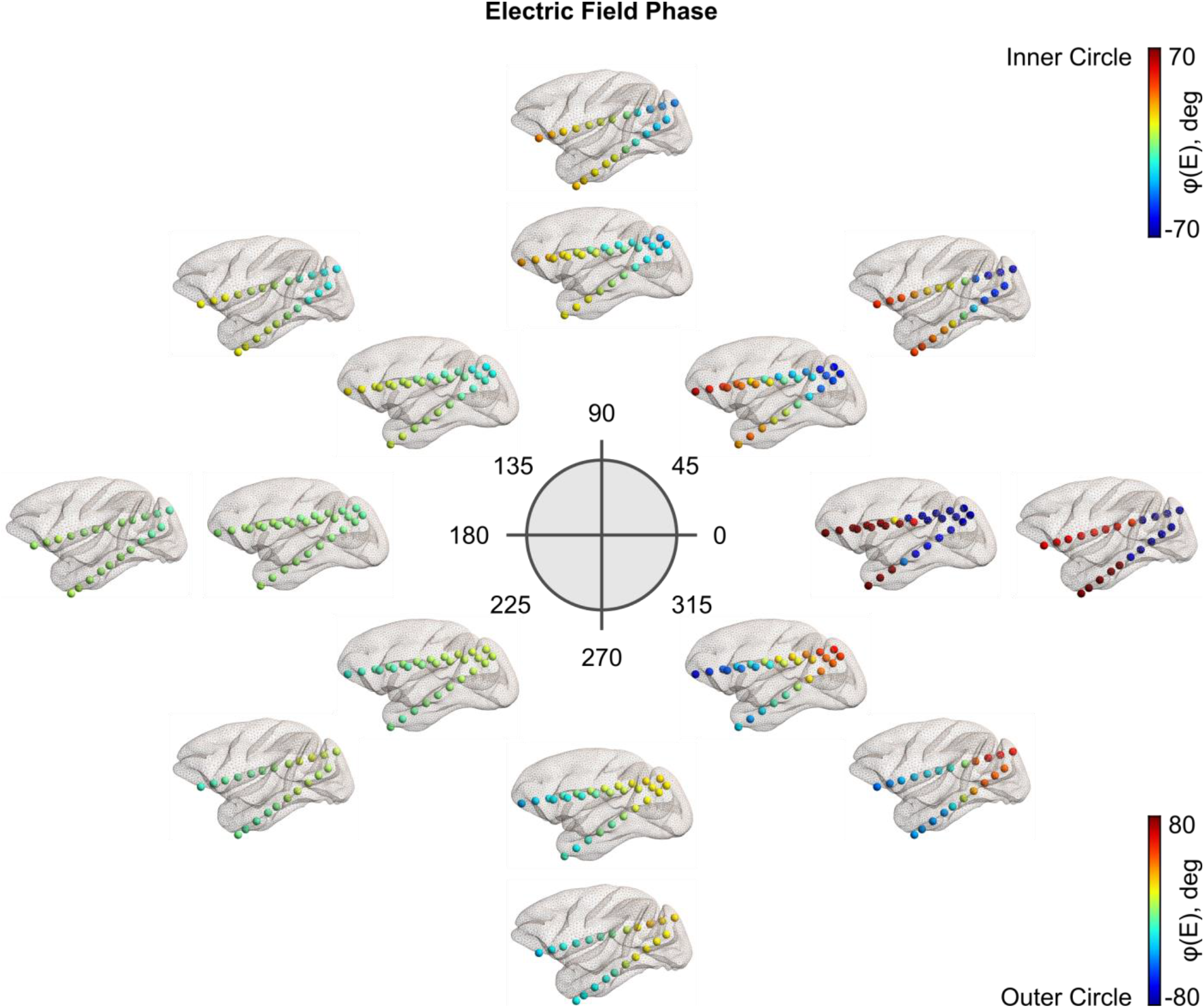
Electric field phases in the brain during TES. The phase difference between the anterior and posterior stimulation electrodes is indicated in the middle plot. The inner circle of plot corresponds to subject 1, and the outer circle to subject 2. This figure complements Figure 4 in the main paper.

**Supplementary Figure 6.**
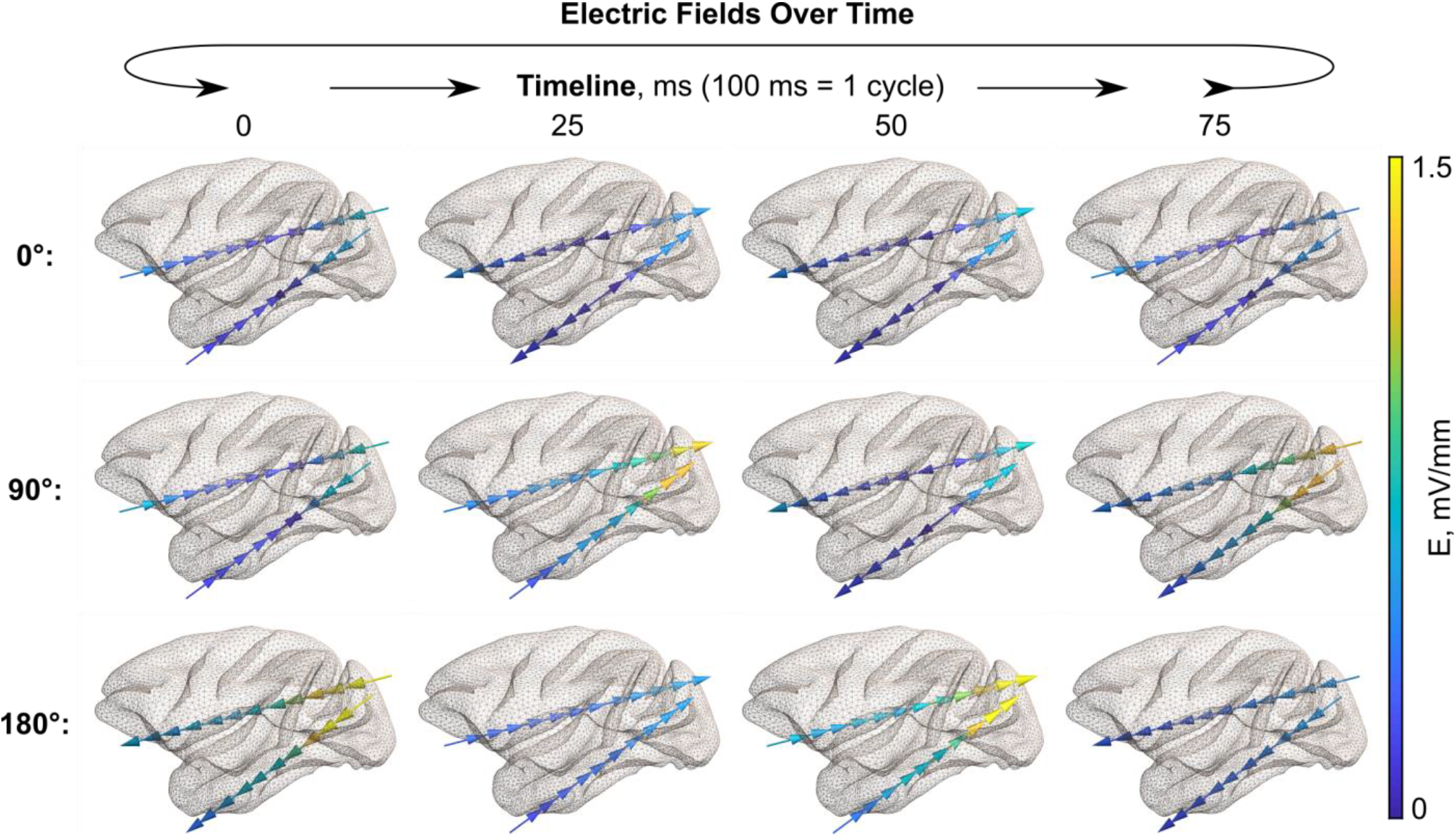
Electric fields in the brain over time during TES for a given stimulation condition for subject 2. The panel depicts the main conditions (0°, 90°, and 180° stimulation phase differences) for subject 2. Arrows indicate the electric field direction, and the color encodes the electric field magnitude. This figure complements the Fig. 5 in the main paper.

**Supplementary Animation 1.**
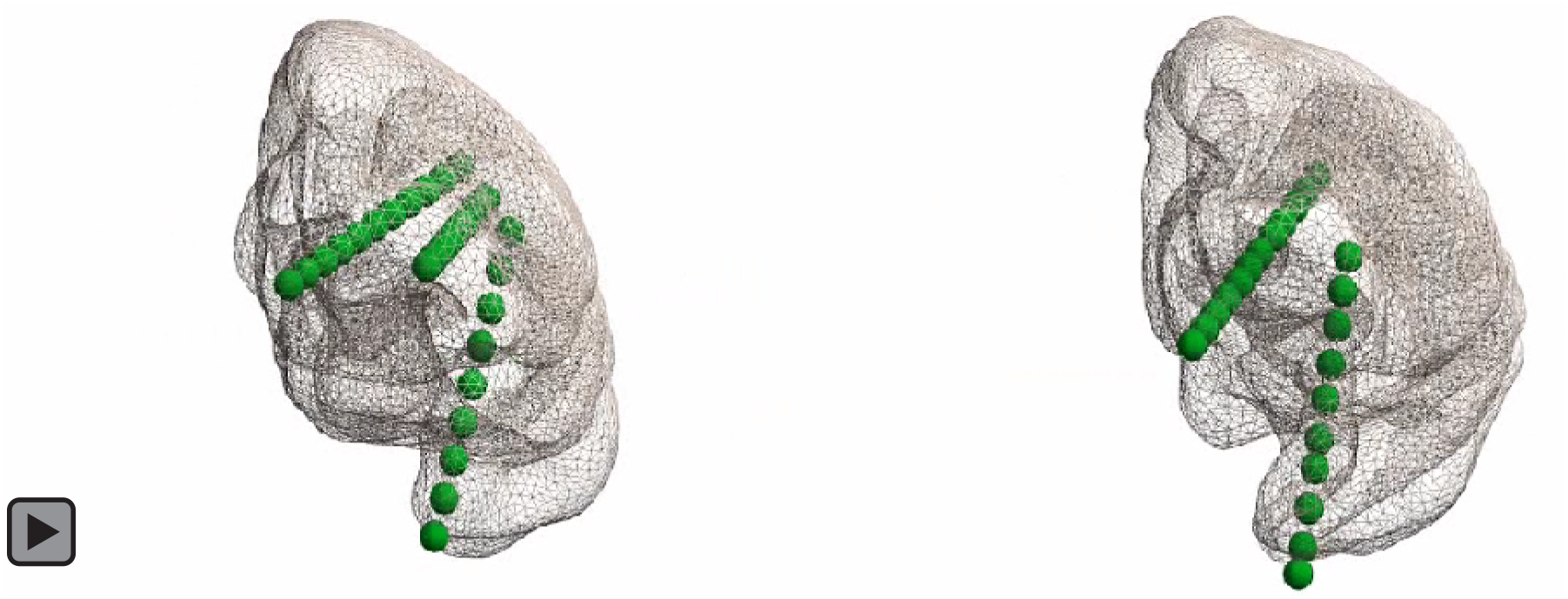
Invasive recording electrodes in the brain of subject 1 (left) and subject 2 (right). Green spheres depict the recording contacts. This animation complements Figure 1 in the main paper.

**Supplementary Animation 2.**
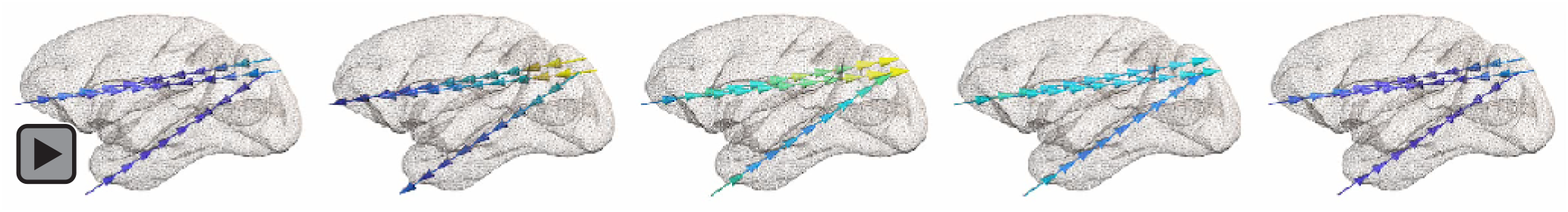
TES electric fields in the brain over time for subject 1. The stimulation conditions (i.e. the current phase differences between the anterior and posterior stimulation electrodes) are equal to 0°, 90°, 180°, 270° and 360° from left to right. The electric field magnitude is color-coded from dark blue (0 mV/mm) to bright yellow (0.8 mV/mm per 1 mA). The arrow directions indicate the electric field direction. This animation complements Figure 5 in the main paper.

**Supplementary Animation 3.**
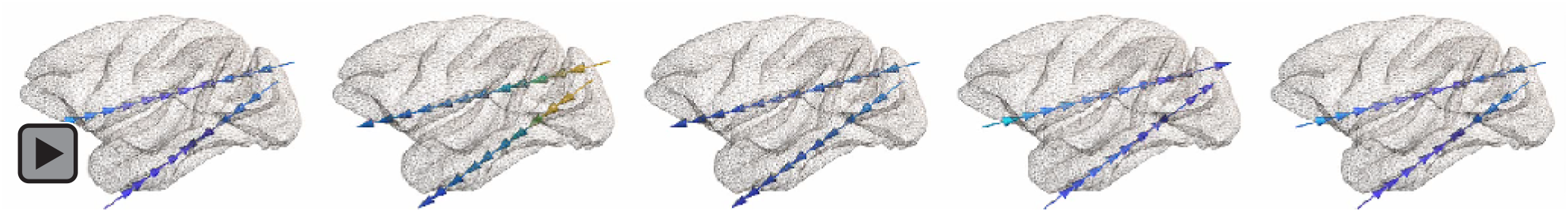
TES electric fields in the brain over time for subject 2. The stimulation conditions (i.e. the current phase differences between the anterior and posterior stimulation electrodes) are equal to 0°, 90°, 180°, 270° and 360° from left to right. The electric field magnitude is color-coded from dark blue (0 mV/mm) to bright yellow (1.2 mV/mm per 1 mA). The arrow directions indicate the electric field direction. This animation complements Figure 5 in the main paper.

**Supplementary Animation 4.**
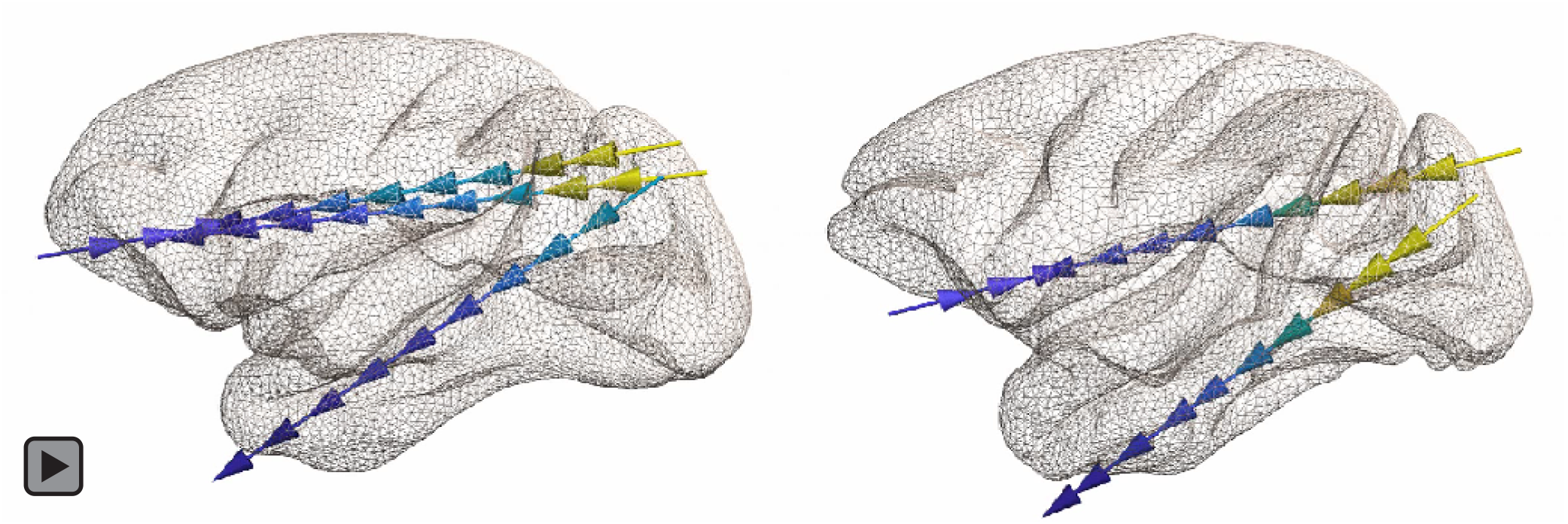
Traveling wave stimulation with the local electric field maximum moving over time. Shown is the 45° stimulation condition in subject 1 (left) and 2 (right). Absolute normalized electric field intensity is color-coded from dark blue (relative minimum) to bright yellow (relative maximum). This animation complements Figure 6 in the main paper.

## REFERENCES

1. Buzsáki, G., Anastassiou, C. A. & Koch, C. The origin of extracellular fields and currents — EEG, ECoG, LFP and spikes. Nat. Rev. Neurosci. 13, 407–420 (2012).

2. Siegel, M., Donner, T. H. & Engel, A. K. Spectral fingerprints of large-scale neuronal interactions. Nat. Rev. Neurosci. 13, 121–134 (2012).

3. Canolty, R. T. & Knight, R. T. The functional role of cross-frequency coupling. Trends Cogn. Sci.14, 506–515 (2010).

4. Schroeder, C. E. & Lakatos, P. Low-frequency neuronal oscillations as instruments of sensory selection. Trends Neurosci. 32, 9–18 (2009).

5. Buzsáki, G., Logothetis, N. & Singer, W. Scaling Brain Size, Keeping Timing: Evolutionary Preservation of Brain Rhythms. Neuron 80, 751–764 (2013).

6. Voytek, B. & Knight, R. T. Dynamic Network Communication as a Unifying Neural Basis for Cognition, Development, Aging, and Disease. Biol. Psychiatry 77, 1089–1097 (2015).

7. Uhlhaas, P. J. & Singer, W. Abnormal neural oscillations and synchrony in schizophrenia. Nat. Rev. Neurosci. 11, 100–113 (2010).

8. Bassett, D. S. & Sporns, O. Network neuroscience. Nat. Neurosci. 20, 353–364 (2017).

9. Anastassiou, C. A. & Koch, C. Ephaptic coupling to endogenous electric field activity: why bother? Curr. Opin. Neurobiol. 31, 95–103 (2015).

10. Polanía, R., Nitsche, M. A. & Ruff, C. C. Studying and modifying brain function with non-invasive brain stimulation. Nat. Neurosci. 21, 174–187 (2018).

11. Yavari, F., Jamil, A., Mosayebi Samani, M., Vidor, L. P. & Nitsche, M. A. Basic and functional effects of transcranial Electrical Stimulation (tES)—An introduction. Neurosci. Biobehav. Rev. 85, 81–92 (2018).

12. Antal, A. et al. Low intensity transcranial electric stimulation: Safety, ethical, legal regulatory and application guidelines. Clin. Neurophysiol. 128, 1774–1809 (2017).

13. Thut, G. et al. Guiding transcranial brain stimulation by EEG/MEG to interact with ongoing brain activity and associated functions: A position paper. Clin. Neurophysiol. 128, 843–857 (2017).

14. Turi, Z., Alekseichuk, I. & Paulus, W. On ways to overcome the magical capacity limit of working memory. PLOS Biol. 16, e2005867 (2018).

15. Polanía, R., Nitsche, M. A., Korman, C., Batsikadze, G. & Paulus, W. The Importance of Timing in Segregated Theta Phase-Coupling for Cognitive Performance. Curr. Biol. 22, 1314–1318 (2012).

16. Helfrich, R. F. et al. Selective Modulation of Interhemispheric Functional Connectivity by HD-tACS Shapes Perception. PLoS Biol. 12, e1002031 (2014).

17. Alekseichuk, I., Turi, Z., Amador de Lara, G., Antal, A. & Paulus, W. Spatial Working Memory in Humans Depends on Theta and High Gamma Synchronization in the Prefrontal Cortex. Curr. Biol. 26, 1513–1521 (2016).

18. Violante, I. R. et al. Externally induced frontoparietal synchronization modulates network dynamics and enhances working memory performance. Elife 6, e22001 (2017).

19. Polanía, R., Moisa, M., Opitz, A., Grueschow, M. & Ruff, C. C. The precision of value-based choices depends causally on fronto-parietal phase coupling. Nat. Commun. 6, 8090 (2015).

20. Bächinger, M. et al. Concurrent tACS-fMRI Reveals Causal Influence of Power Synchronized Neural Activity on Resting State fMRI Connectivity. J. Neurosci. 37, 4766–4777 (2017).

21. Tseng, P., Iu, K.-C. & Juan, C.-H. The critical role of phase difference in theta oscillation between bilateral parietal cortices for visuospatial working memory. Sci. Rep. 8, 349 (2018).

22. Strüber, D., Rach, S., Trautmann-Lengsfeld, S. A., Engel, A. K. & Herrmann, C. S. Antiphasic 40 Hz Oscillatory Current Stimulation Affects Bistable Motion Perception. Brain Topogr. 27, 158–171 (2014).

23. Alekseichuk, I., Pabel, S. C., Antal, A. & Paulus, W. Intrahemispheric theta rhythm desynchronization impairs working memory. Restor. Neurol. Neurosci. 35, 147–158 (2017).

24. Fries, P. Rhythms for Cognition: Communication through Coherence. Neuron 88, 220–235 (2015).

25. Fell, J. & Axmacher, N. The role of phase synchronization in memory processes. Nat. Rev. Neurosci. 12, 105–118 (2011).

26. Hanslmayr, S., Staresina, B. P. & Bowman, H. Oscillations and Episodic Memory: Addressing the Synchronization/Desynchronization Conundrum. Trends Neurosci. 39, 16–25 (2016).

27. Thut, G., Miniussi, C. & Gross, J. The Functional Importance of Rhythmic Activity in the Brain. Curr. Biol. 22, R658–R663 (2012).

28. Saturnino, G. B., Madsen, K.H., Siebner, H. R. & Thielscher, A. How to target inter-regional phase synchronization with dual-site Transcranial Alternating Current Stimulation. Neuroimage 163, 68–80 (2017).

29. Opitz, A. et al. Spatiotemporal structure of intracranial electric fields induced by transcranial electric stimulation in humans and nonhuman primates. Sci. Rep. 6, 31236 (2016).

30. Opitz, A., Falchier, A., Linn, G. S., Milham, M. P. & Schroeder, C. E. Limitations of ex vivo measurements for in vivo neuroscience. Proc. Natl. Acad. Sci. 114, 5243–5246 (2017).

31. Krause, M. R. et al. Transcranial Direct Current Stimulation Facilitates Associative Learning and Alters Functional Connectivity in the Primate Brain. Curr. Biol. 27, 3086–3096.e3 (2017).

32. Kar, K., Duijnhouwer, J. & Krekelberg, B. Transcranial Alternating Current Stimulation Attenuates Neuronal Adaptation. J. Neurosci. 37, 2325–2335 (2017).

33. Oostenveld, R., Fries, P., Maris, E. & Schoffelen, J.-M. FieldTrip: Open Source Software for Advanced Analysis of MEG, EEG, and Invasive Electrophysiological Data. Comput. Intell. Neurosci. 2011, 156869 (2011).

34. Geuzaine, C. & Remacle, J.-F. Gmsh: A 3-D finite element mesh generator with built-in pre- and post-processing facilities. Int. J. Numer. Methods Eng. 79, 1309–1331 (2009).

35. Radman, T., Ramos, R. L., Brumberg, J. C. & Bikson, M. Role of cortical cell type and morphology in subthreshold and suprathreshold uniform electric field stimulation in vitro. Brain Stimul. 2, 215–28, 228.e1–3 (2009).

36. Rawji, V. et al. tDCS changes in motor excitability are specific to orientation of current flow. Brain Stimul. 11, 289–298 (2018).

37. Rahman, A. et al. Cellular effects of acute direct current stimulation: somatic and synaptic terminal effects. J Physiol 591, 2563–2578 (2013).

38. Noury, N. & Siegel, M. Phase properties of transcranial electrical stimulation artifacts in electrophysiological recordings. Neuroimage 158, 406–416 (2017).

39. Muller, L., Chavane, F., Reynolds, J. & Sejnowski, T. J. Cortical travelling waves: mechanisms and computational principles. Nat. Rev. Neurosci. 19, 255–268 (2018).

40. Wu, J.-Y., Xiaoying Huang & Chuan Zhang. Propagating Waves of Activity in the Neocortex: What They Are, What They Do. Neurosci. 14, 487–502 (2008).

41. Patel, J., Fujisawa, S., Berényi, A., Royer, S. & Buzsáki, G. Traveling Theta Waves along the Entire Septotemporal Axis of the Hippocampus. Neuron 75, 410–417 (2012).

42. Rubino, D., Robbins, K. A. & Hatsopoulos, N. G. Propagating waves mediate information transfer in the motor cortex. Nat. Neurosci. 9, 1549–1557 (2006).

43. Besserve, M., Lowe, S. C., Logothetis, N. K., Schölkopf, B. & Panzeri, S. Shifts of Gamma Phase across Primary Visual Cortical Sites Reflect Dynamic Stimulus-Modulated Information Transfer. PLOS Biol. 13, e1002257 (2015).

44. Massimini, M., Huber, R., Ferrarelli, F., Hill, S. & Tononi, G. The Sleep Slow Oscillation as a Traveling Wave. J. Neurosci. 24, 6862–6870 (2004).

45. Bahramisharif, A. et al. Propagating Neocortical Gamma Bursts Are Coordinated by Traveling Alpha Waves. J. Neurosci. 33, 18849–18854 (2013).

46. Huang, Y. et al. Measurements and models of electric fields in the in vivo human brain during transcranial electric stimulation. Elife 6, e18834 (2017).

47. Windhoff, M., Opitz, A. & Thielscher, A. Electric field calculations in brain stimulation based on finite elements: An optimized processing pipeline for the generation and usage of accurate individual head models. Hum. Brain Mapp. 34, 923–935 (2013).

48. Reato, D., Rahman, A., Bikson, M. & Parra, L. C. Low-Intensity Electrical Stimulation Affects Network Dynamics by Modulating Population Rate and Spike Timing. J. Neurosci. 30, 15067–15079 (2010).

49. Terzuolo, C. A. & Bullock, T. H. Measurement of imposed voltage gradient adequate to modulate neuronal firing. Proc. Natl. Acad. Sci. 42, 687–694 (1956).

50. Fröhlich, F. & McCormick, D. A. Endogenous Electric Fields May Guide Neocortical Network Activity. Neuron 67, 129–143 (2010).

51. Anastassiou, C. A., Perin, R., Markram, H. & Koch, C. Ephaptic coupling of cortical neurons. Nat. Neurosci. 14, 217–223 (2011).

52. Vöröslakos, M. et al. Direct effects of transcranial electric stimulation on brain circuits in rats and humans. Nat. Commun. 9, 483 (2018).

53. Opitz, A., Paulus, W., Will, S., Antunes, A. & Thielscher, A. Determinants of the electric field during transcranial direct current stimulation. Neuroimage 109, 140–150 (2015).

54. Miranda, P. C., Mekonnen, A., Salvador, R. & Ruffini, G. The electric field in the cortex during transcranial current stimulation. Neuroimage 70, 48–58 (2013).

55. Laakso, I. et al. Electric fields of motor and frontal tDCS in a standard brain space: A computer simulation study. Neuroimage 137, 140–151 (2016).

